# hC9ORF78 localizes to kinetochores and is required for proper chromosome segregation

**DOI:** 10.1101/2021.02.03.429653

**Authors:** Radhika Koranne, Kayla M. Brown, Hannah E. Vandenbroek, William R. Taylor

## Abstract

C9ORF78 is a poorly characterized protein found in diverse eukaryotes. Previous work indicated overexpression of hC9ORF78 (aka HCA59) in malignant tissues indicating a possible involvement in growth regulatory pathways. Additional studies in fission yeast and humans uncover a potential function in regulating the spliceosome. In studies of GFP-tagged hC9ORF78 we observed a dramatic reduction in protein abundance in cells grown to confluence and/or deprived of serum growth factors. Serum stimulation induced synchronous re-expression of the protein in HeLa cells. This effect was also observed with the endogenous protein. Overexpressing either E2F1 or N-Myc resulted in elevated hC9ORF78 expression potentially explaining the serum-dependent upregulation of the protein. Immunofluorescence analysis indicates that hC9ORF78 localizes to nuclei in interphase but does not appear to concentrate in speckles as would be expected for a splicing protein. Surprisingly, a subpopulation of hC9ORF78 co-localizes with ACA, Mad1 and Hec1 in mitotic cells suggesting that this protein may associate with kinetochores or centromeres. Furthermore, knocking-down hC9ORF78 caused mis-alignment of chromosomes in mitosis. These studies uncover novel mitotic function and subcellular localization of cancer antigen hC9ORF78.

**SUMMARY STATEMENT:** hC9ORF78 regulates chromosome segregation.

## Introduction

Multiple mechanisms ensure high fidelity of chromosome segregation to create viable daughter cells. Since attachment of chromosomes to the spindle involves random search and capture, checkpoint and surveillance mechanisms exist to correct errors in attachment [1–4]. Errors that go uncorrected before cell division result in aneuploidy or cell death. The mitotic checkpoint coordinates chromosome segregation by inhibiting anaphase onset until all chromosomes attain bipolar attachment to the spindle [4]. Unattached kinetochores recruit checkpoint proteins Mad1 and Mad2 which catalyze the formation of Mitotic Checkpoint Complex (MCC) an inhibitor of the Anaphase Promoting Complex/Cyclosome (APC/C). MCC is composed of Mad2, Cdc20, Bub3 and BubR1 and is generated as long as unattached kinetochores persist. When all kinetochores attach to the spindle, loss of MCC allows activation of APC/C, an E3 ubiquitin ligase that targets Cyclin B and Securin for destruction allowing exit from mitosis [5]. Certain cancer types show clear defects in MCC components which include mutations and abnormal expression levels [6–9]. These defects weaken the mitotic checkpoint resulting in abnormal chromosome segregation and aneuploidy. While the mitotic checkpoint detects unattached kinetochores, more subtle attachment defects (ex. both kinetochores attached to same spindle pole) are corrected by the chromosomal passenger complex (CPC) surveillance system [1]. CPC, composed of Aurora B, Borealin, Survivin, and INCENP localizes to the inner centromere and kinetochore and destabilizes defective attachments thereby triggering the mitotic checkpoint until attachment defects are corrected [10–12]. At least 100 proteins localized to kinetochores coordinate attachment to the spindle and ensure operation of the mitotic checkpoint and CPC surveillance system [13]. Many details relevant to the function of the kinetochore are unknown.

C9ORF78 (also known as HCA59 and TLS1) is found in diverse eukaryotes including yeast, plants, animals, and various protist orders indicating a relatively ancient origin (NCBI-blast). Human C9ORF78 (hC9ORF78) is a nuclear protein containing 289 amino acids with no obvious functional domains. Published reports focused on this protein are minimal. In one study, C9ORF78 was identified as a human cancer antigen and was given the alias HCA59. A putative yeast orthologue, TLS1 regulates splicing and while the human protein was found transiently associated with the spliceosome, it is not known if hC9ORF78 regulates splicing. Here we identify a subpopulation of hC9ORF78 localized to kinetochores. Suppressing hC9ORF78 expression causes defects in chromosome segregation. Consistent with a cell cycle role, we also find that hC9ORF78 is a serum-induced protein that is also induced by E2F1 and Myc. These studies uncover a novel mitotic function of a poorly characterized human cancer antigen.

## Results

### Regulation of hC9ORF78 expression

In a previous study, hC9ORF78 was identified as a cancer antigen and although it was reported to be overexpressed in cancer cell lines, these data were not shown. In a limited survey, we observed that hC9ORF78 was significantly overexpressed in breast cancer cell lines MDA MB 231, MDA MB 468, lung cancer cell line NCI H522, colon cancer HCT116 and cervical cancer HeLaM compared to normal retinal epithelial cells (RPE) and normal WI38 fibroblasts (1.2-20X fold range, pvalue <0.05, Supplemental figure 1). A genome-wide CRISPR screen suggested that hC9ORF78 may be required for efficient growth of cancer cells in 3D culture [14]. In a preliminary screen, we observed smaller colonies in HCT116 cultures transiently transfected with hC9ORF8 gRNAs. Also, we were unable to obtain clones of HeLaM cells lacking expression of hC9ORF78 using CRISPR knock-out, potentially consistent with a role in proliferation.

To further analyze hC9ORF78 expression, we created an N-terminal GFP fusion and established stable clones in HeLa cells. We observed that GFP-hC9ORF78 fluorescence was diminished in crowded cultures. Addition of fresh medium containing serum caused a gradual but consistent increase in GFP-hC9ORF78 fluorescence (Fig. 1A). Image analysis of individual cells indicated maximal fluorescence at ~14 hours post medium change (Fig. 1B). Fluorescence imaging of interphase cells indicated that most GFP-hC9ORF78 was localized to the nucleus (Fig. 1). Similar results were obtained using immunofluorescence to detect the endogenous protein (our unpublished data). Next, we used immunofluorescence to measure the behavior of endogenous hC9ORF78 in response to serum stimulation. Image analysis indicated low levels of staining in serum starved cells and an increase after serum induction (Fig. 1C). Serum induction was more dramatic in normal RPE cells compared to HCT116 colon cancer cells. Serum induction of hC9ORF78 broadly paralleled the elevation of Ki67 staining to mark proliferating cells. The two antigens were correlated in RPE (R^2^=0.69) but not HCT116 cells (R^2^=0.26) after addition of 20% serum.

**Figure 1.**
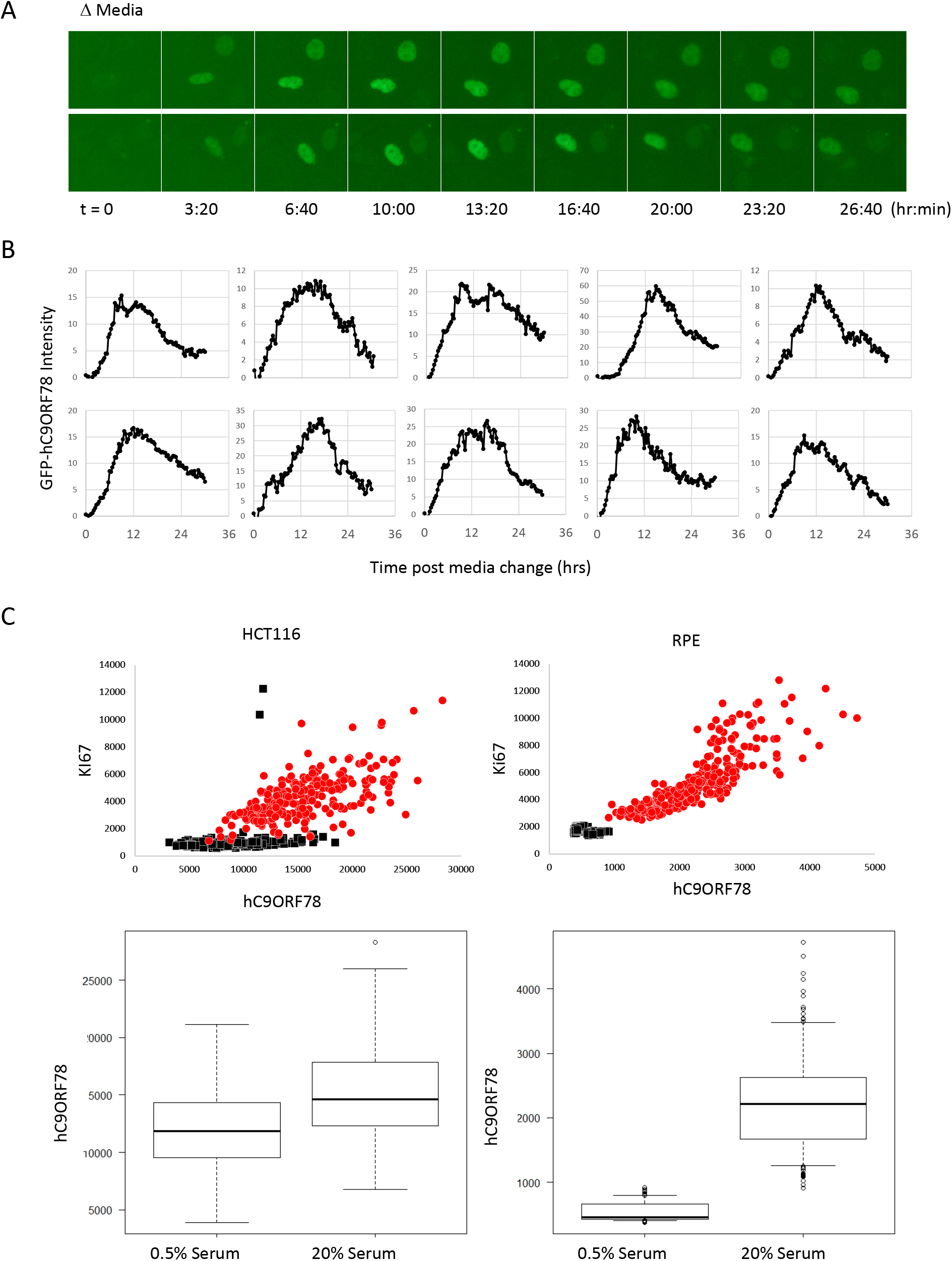
Expression of hC9ORF78. GFP-hC9ORF78 fluorescence in response to media change. HeLa cells stably expressing GFP-hC9ORF78 were grown to confluence and analyzed by time-lapse fluorescence imaging. Imaging was carried out at 37°C using a sealed flask containing 10%CO2. (A) Examples of cells showing increased fluorescence after media change. (B) Fluorescence intensity in 10 cells was quantified to determine the kinetics of recovery. (C) Endogenous hC9ORF78 expression in response to serum stimulation. HCT116 or RPE cells were analyzed by quantitative immunofluorescence. Cells were grown for 3 days in low (0.5%) serum. New media containing either 0.5% (black dots) or 20% serum (red dots) was added and cells were fixed 48 hours later. Immunofluorescence staining was carried out using antibodies to hC9ORF78 and the proliferation marker Ki67. Images were captured, and nuclear areas defined using ImageJ based on Hoechst staining. hC9ORF78 and Ki67 intensity was then determined in the nuclear area. In the top panels, each dot represents one cell. Whisker plots are shown in the bottom panels.

Induction of endogenous hC9ORF78 by serum was also observed by western blotting (Fig. 2A). Genome-wide ChIP sequencing suggests that hC9ORF78 may be regulated by E2F, Myc and Elf1 [15, 16]. Overexpressing E2F1 using a recombinant adenovirus caused upregulation of hC9ORF78 in serum starved normal WI38 cells (Fig. 2A, B, C). Also, Rat embryonic fibroblasts over-expressing N-Myc showed higher levels of hC9ORF78 as compared to wild type (Fig. 2D, E)[17]. Expression of a dominant-negative mutant of p53 had no significant effect on hC9ORF78 expression (Fig. 2 D and E). Thus, E2F1 and Myc regulation at least partly explains upregulation of hC9ORF78 by serum. However, transcriptional induction is unlikely to explain elevation of GFP-hC9ORF78 which is transcriptionally driven by a CMV promoter in the transfected plasmid (Fig. 1).

**Figure 2.**
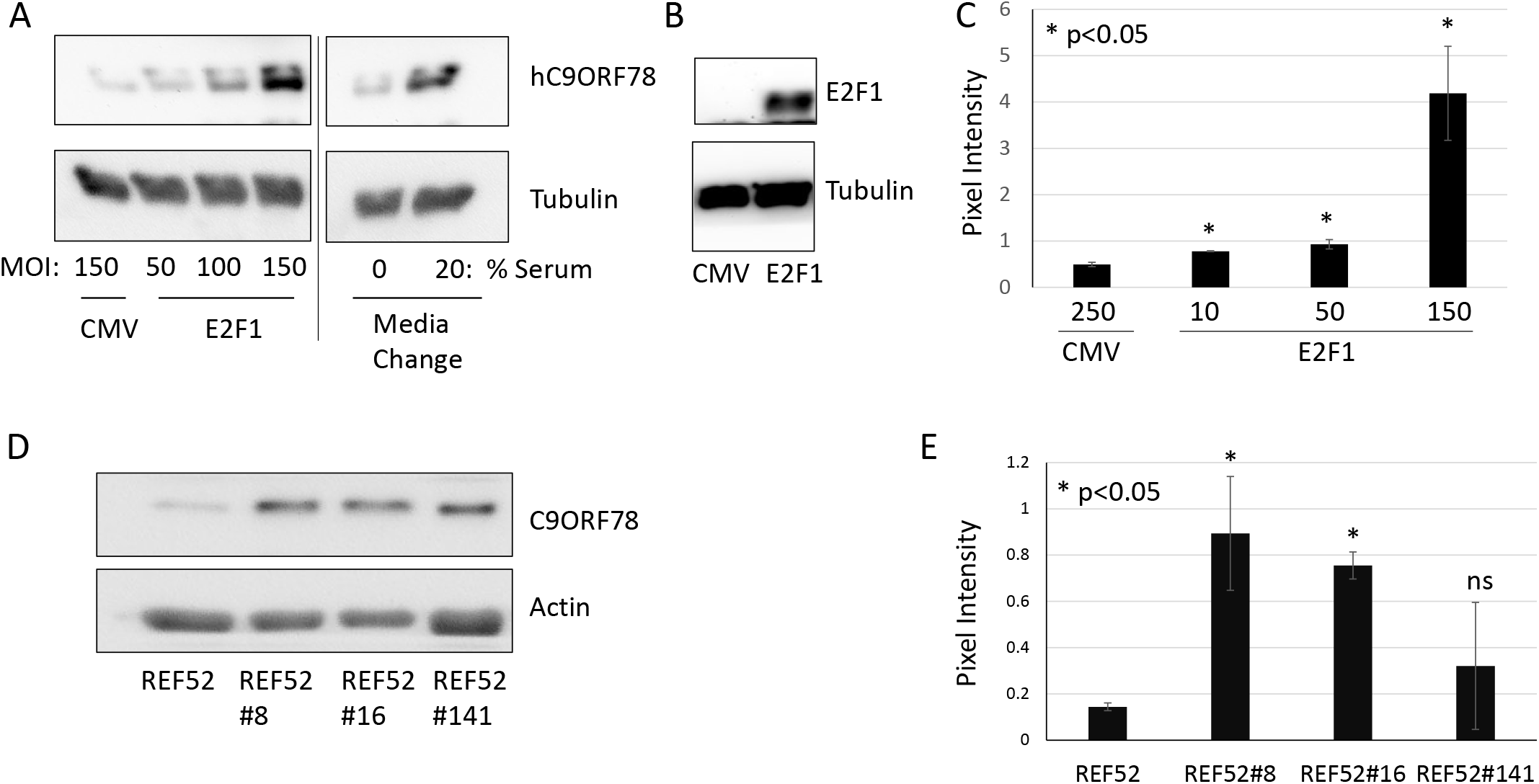
Regulation of hC9ORF78 by E2F1 and N-Myc. Induction of hC9ORF78 by serum, E2F1 and N-Myc. Western blotting was used to measure endogenous hC9ORF78 in either WI-38 cells or nontransformed rat cell line REF52. (A) WI-38 cells were either induced by serum stimulation or infected with recombinant adenoviruses at the indicated multiplicities of infection (MOI). Adenovirus with no transgene (CMV) was used as a control for virus expressing E2F1. E2F1 overexpression is shown in (B). (C) hC9ORF78 levels detected by western blotting were quantified in three independent experiments. *p< 0.05 versus CMV (D and E) Parental REF52 cells or clones overexpressing either N-Myc or the C141Y dominant-negative p53 mutant were analyzed by western blotting. Example of western analysis is shown in (D) and quantitation of triplicate experiments is shown in (E). Average and standard deviations are shown. *p< 0.05 versus REF52.

### Localization of hC9ORF78 in mitosis

We analyzed asynchronously growing HeLa cells using immunofluorescence with antibodies to detect the endogenous protein. Confocal imaging revealed dispersed cytoplasmic staining during mitosis but also foci that coincided with anti-centromere (ACA) antibody staining (Fig. 5, 6). Cells were pre-extracted with 0.1% triton X-100 for 0.5 minutes before fixing with formaldehyde followed by standard immunofluorescence staining. Cells were co-stained with antibodies to either Ndc80 or Mad1 to detect kinetochores. Under these conditions, hC9ORF78 colocalized with Ndc80 and Mad1 indicating that this protein localizes to the kinetochore (Fig. 3A, B and C). Foci of hC9ORF78 staining that do not colocalize with centromere/kinetochore markers were also observed (Fig. 3C). Next, we quantified centromere-proximal hC9ORF78 pixel intensity by creating regions of interest based on ACA staining and collecting image data from both antigen channels. Most kinetochores showed a hC9ORF78/ACA pixel intensity ratio of ~0.58. This ratio was significantly decreased to ~0.22 in HeLa cells transfected with RNAi targeting hC9ORF78 indicating that the centromere-proximal foci detected by the hC9ORF78 antibodies are specific (p<0.05; Supplemental Fig. 2).

**Figure 3.**
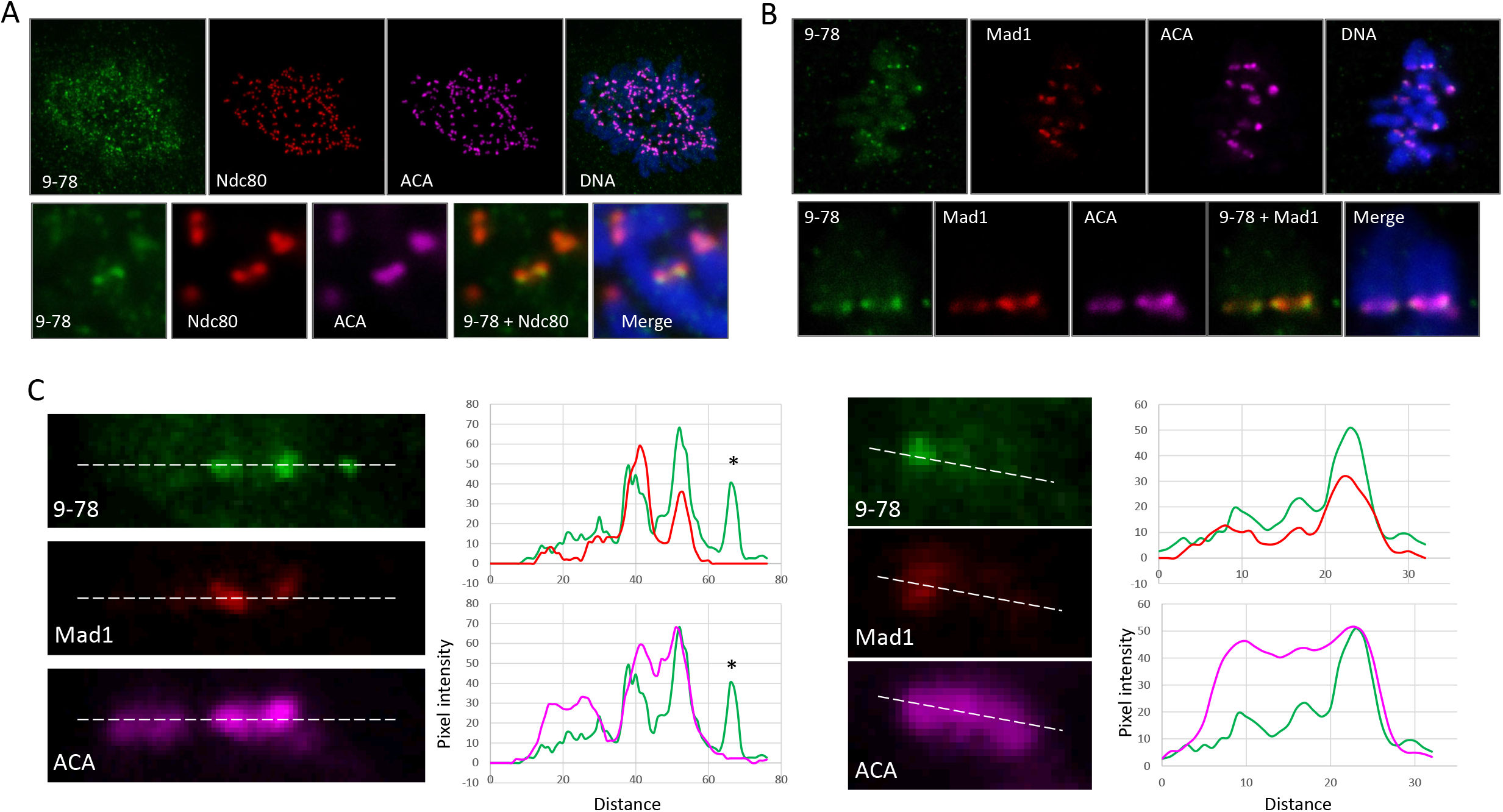
Kinetochore localization of hC9ORF78. Asynchronously growing HeLa cells were pre-extracted with triton X-100, fixed and analyzed by immunofluorescence using a polyclonal hC9ORF78 antiserum (9-78: Bethyl) and anticentromere antibodies (ACA: Antibodies INC). Alexafluor-conjugated secondary antibodies were from Invitrogen. DNA was visualized with Hoechst 33343. Confocal microscopy was used to detect fluorescent signals. Single z-planes are shown. Cells were also stained with either (A) Ndc80/Hec1 (B) or Mad1 and ACA. (C) Line scans showing hC9ORF78 overlap with Mad1. *indicates presence of non-kinetochore hC9ORF78 foci.

### Chromosome mis-segregation upon hC9ORF78 knockdown

Initial observations suggested that HeLa cells transfected with RNAi targeting hC9ORF78 exhibited chromosome segregation defects. To quantify this effect, we repeated the transfection and also included an additional cancer line HCT116. Cells were fixed and stained with antibodies to ACA, and tubulin to visualize the mitotic spindle. Sample identities were shielded so that chromosome mis-segregation could be assessed in a blinded manner. Mis-segregation was defined by the presence of chromosomes that had not congressed to the metaphase plate. In multiple independent experiments, hC9ORF78 knockdown cells showed an increase in chromosome mis-segregation (Fig. 4).

**Figure 4.**
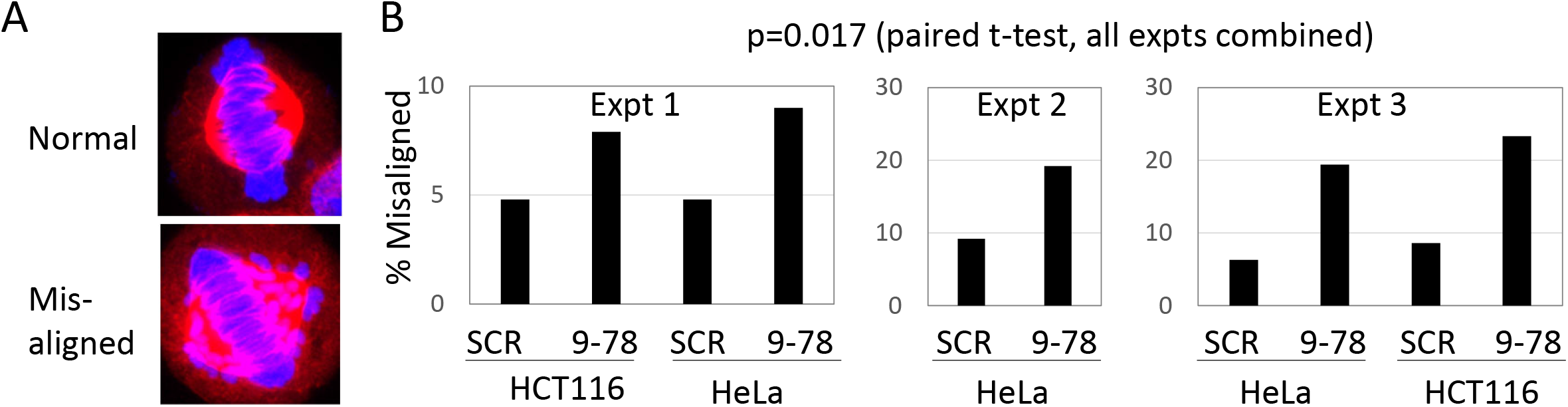
Chromosome mis-segregation upon hC9ORF78 knockdown. Chromosome mis-segregation upon hC9ORF78 knockdown. HCT116 or HeLa cells were transfected with RNAi targeting hC9ORF78 (9-78) or non-targeting control (SCR). Chromosome segregation was quantified in a blinded manner using samples analyzed by immunofluorescence staining. (A) Example of a HeLa cell with mis-aligned chromosomes. (B) Quantification of mis-alignment phenotype.

## Discussion

C9ORF78 is a poorly characterized protein with an ancient evolutionary origin. Orthologues exists in most of the eukaryotic kingdoms (NCI-BLAST). Limited studies suggest a potential role in splicing, regulation of ubiquitination and association with cancer depending on the species under study. hC9ORF78 was previously named hepatocellular antigen (HCA)59 based on the finding of antibodies in patients with that particular cancer [18]. In that study, HCA59 was reported to be overexpressed in several cancer cell lines, a finding we have also corroborated. Our failed attempts to generate an hC9ORF78 knock-out HeLa clone implies its role in cancer cell proliferation. Several genome wide CRISPR-Cas9 screens suggest that hC9ORF78 could be involved in cell proliferation and oncogenesis[14, 19, 20]. A potential plant orthologue, CSU2, affects photomorphogenesis by regulating the E3 ligase COP1 [21]. Whether hC9ORF78 regulates the ubiquitin system has not been directly examined. The putative orthologue in fission yeast, named TLS1, regulates the splicing of rap1+ and poz1+, components of the shelterin complex that protects telomeres [22]. Also, hC9ORF78 was identified as a substoichiometric component of complex C of the spliceosome, however, the functional relevance of this association was not investigated [23]. Using GFP-trap pull-down in cells expression GFP-hC9ORF78, we confirmed that hC9ORF78 associates with the complex C protein Pre-mRNA processing Factor 8 (Supplemental figure 3A). However, we observed that hC9ORF78 does not significantly affect the expression levels of shelterin components as observed for the fission yeast orthologue (Supplemental Fig. 3B, C, D). Therefore, if hC9ORF78 regulates splicing, its targets are yet to be defined. Further studies to determine function of hC9ORF78 in splicing machinery are needed.

Genes known or likely to be involved in proliferation show distinct upregulation after serum stimulation in quiescent fibroblast cells. Upregulation of genes involved in cell cycle progression such as Cyclins, CDKs, DNA replication factors and several transcription factors after serum stimulation is well documented [24]. Our studies show that hC9ORF78 is a serum-induced protein which is as least partly explained by E2F1 and Myc-dependent regulation. Immunofluorescence analysis uncovered a population of hC9ORF78 proximal to kinetochores. Knocking down expression of hC9ORF78 created errors in chromosome segregation indicating a novel mitotic role of this protein. Future studies are focused on how hC9ORF78 is recruited to kinetochores and the mechanism by which hC9ORF78 regulates chromosome segregation. There are numerous reports linking various aspects of RNA metabolism to kinetochore function. For example, transcription of ncRNAs at the centromere is essential for proper kinetochore assembly and function [25]. In one study, splicing of these ncRNAs was also implicated in kinetochore function [26]. Indeed, splicing protein PRP4 was found to localize to kinetochores to regulate the mitotic checkpoint [27]. Therefore, it is possible that hC9ORF78 localizes to kinetochores to regulate splicing of ncRNAs. Alternatively, the role of hC9ORF78 at kinetochores maybe independent of its interaction with complex C of the spliceosome. In summary, we provide evidence for a novel role of hC9ORF78 in chromosome segregation and uncover a kinetochore-localized population of the protein that may fulfill this role.

## Materials and Methods

### Cell Lines and Culture Conditions

Cell lines were cultured at 37°C in a humidified atmosphere containing 10% CO2 in Dulbecco’s Modified Eagle’s Medium (Mediatech, Inc.) supplemented with 10% fetal bovine serum (Sigma) or calf serum (Atlanta Biologicals) and Penicillin/Streptomycin 100U/ml. Cell types used; WI-38 (human embryonic lung fibroblast), HeLaM (a subclone of HeLa [10]), HCT116, hTERT RPE, MDA-MB-231, MDA-MB-468, H522, HEK239 and REF-52.

### Plasmids and RNAi transfections

Entry clones were prepared using hC9ORF78 cDNA (DNAsu) and pENTR directional TOPO cloning kit (Invitrogen). The entry clone was confirmed by sequencing. To GFP tag hC9ORF78 at N-terminus, the entry clone was subcloned into dECE-GFP. To knock-in GFP at the endogenous locus of hC9ORF78, we used MMEJ assisted gene knock in using CRISPR-cas9 and PITCh system[28]. Briefly to prepare repair template 20 nucleotide long microhomology region of hC9ORF78 was cloned flanking EGFP-2A-Puro region in PITCh vector (upstream microhomology region- 5’ CCGCGTTACATAGCATCGTACGCGTACGTGTTTGGACTGATGACTATCATTATGAGAAGTTCAAGAAAATGAATCCCCCCGGATCCATGGTGAGCAAGGG 3’, downstream homology region- 5’ ACGCGTACGTGTTTGGAAGGCGATATTTACATCCCACTCTGCACAACTCAGTACCGTCAGGCACCGGGCTTGCG). gRNAs were designed against hC9ORF78 last exon (Top- 5’ CACCGAGAAGTTCAAGAAAATGAAT 3’, Bottom- 5’ AAACATTCATTTTCTTGAACTTCTC 3’). These were cloned into pSpCas9BB-2A-Puro (Addgene). To knock-down hC9ORF78, 75nM ON-TARGET plus siRNA SMARTpool against hC9ORF78 (020231-02) was transfected in 60% confluent 24 well plate using Lipofectamine 3000 (Invitrogen). Scrambled siRNA was used as a control. Cells were fixed/harvested 3 days post transfection.

### Cloning and generation of stable cell lines

To generate HeLaM cell line stably over-expressing GFP-hC9ORF78, cells were transfected with 1 μg pDECE-GFP-hC9ORF78 and a plasmid providing Hygromycin resistance. An isolated colony expressing GFP and resistant to Hygromycin was used for further experiments. To generate clones with GFP knocked-in at endogenous C terminus of hC9ORF78, HeLa cells were transfected with 3 plasmids: a plasmid encoding hC9ORF78 gRNA, PITCh gRNA and PITCh vector (that contains microhomology regions flanking GFP and Puromycin resistance gene). Colonies grown from single cell were screened for Puromycin resistance and nuclear GFP expression. Successful GFP tagging of hC9ORF78 was confirmed by western blotting.

### Western Blotting

Cells were harvested by scraping and lysed in RIPA buffer [10 mM Tris (pH 7.4), 150 mM NaCl, 1% NP-40, 1% DOC, 0.1% SDS, 1 mg/ml aprotinin, 2 mg/ml leupeptin, 1 mg/ml pepstatin A, 1 mM DTT, 0.1M phenylmethylsulfonyl fluoride, 1 mM sodium fluoride and 1 mM sodium vanadate] for 20 minutes on ice. Insoluble debris was removed by centrifugation at 16,000 g for 20 minutes at 4°C. Equal amounts of protein for each sample (determined using BCA protein assay kit - Pierce) were separated by sodium dodecyl sulfate polyacrylamide gel electrophoresis (SDS-PAGE). Gels were transferred to polyvinylidene difluoride membranes (Millipore), blocked in a solution containing 5% (w/v) non-fat dry milk dissolved in PBST [PBS containing 0.05% (v/v) Tween 20], and probed with antibodies to hC9ORF78 (Atlas antibodies and Bethyl), α-tubulin (Sigma), Actin (Abcam), PRPF8 (Abbexa), GFP(SCBT), TERF2IP (Proteintech), POT1 (Proteintech), E2F1(SCBT) Signals were detected using horse-radish peroxidase conjugated secondary antibodies (Biorad) and enhanced chemiluminescence (Biorad).

### Immunofluorescence

cells were plated on sterilized coverslips. After appropriate treatments/transfections they were pre-extracted with permeabilization buffer [150 mM NaCl, 10 mM Tris (pH 7.7), 0.1% Triton X-100, and 0.1% BSA] for 0.5 min. Then fixed with 2% formaldehyde in phosphate buffered saline (PBS) for 10 min, followed by permeabilization buffer for 9 min. Fixed cells were blocked with PBS containing 0.1% BSA (PBSP) for 1 hr at room temperature. Cells were then stained with primary antibodies. Primary antibody dilutions were determined empirically for each antibody. Antibodies were visualized by incubating samples with Alexa-fluor-conjugated secondary antibodies (Invitrogen) 1:1000 diluted in PBSP. DNA was visualized by staining with Hoechst 33342 (Molecular Probes). Phenotypes were counted in blinded manner. Images were captured on SP8 Leica Confocal microscope or Wide field Olympus microscope. Laser intensities, Z-step size and other image acquisition settings were kept constant for a given experiment.

### Image analysis

For analysis of hC9ORF78 signal post serum stimulation, only one Z plane was captured. Nuclear signal (Hoechst) was used as mask in ImageJ. At least 100 cells from each treatment were measured. Intensity of hC9ORF78 and Ki67 for each nucleus was measured. To measure the hC9ORF78/ACA ratio after RNAi knockdown, mitotic cells were analyzed by immunofluorescence. Images were captured on SP8 Leica confocal microscope, one Z plane was used and ROI were drawn using the ACA channel to define centromeres. Using the same ROI, signal intensities were measured for ACA and hC9ORF78 channels. hC9ORF78/ACA signal ratios were calculated, and frequency distributions shown in Supplemental Figure 2.

### Statistical methods

Statistical analyses and drawing graphs were performed using R programming or MS excel. All results were confirmed by multiple independent experiments, quantified by students t-test. p value<0.05 were considered statistically significant. Quantitation of chromosome mis-alignment was conducted in a blinded manner.

### GFP-Trap

One x 10^6^ cells/10cm dish expressing unfused pGFP or GFP tagged to hC9ORF78 were plated. For pulling down GFP tagged proteins, cells from 5 plates were pooled by scraping and whole cell lysates were collected as mentioned in western blotting. Manufacturer protocol (Chromotek) was used further. Briefly 25μl GFP-Trap MA beads equilibrated in dilution buffer [10mM Tris-Cl/ pH7.5, 150mM NaCl, 0.5mM EDTA, protease inhibitors] were added to whole cell lysates diluted in dilution buffer. Beads were collected using a magnet, washed 3 times in dilution buffer, and analyzed by SDS-PAGE.

### Adenoviral Vectors

One 70% confluent HEK293 cells plate was infected with second-generation recombinant adenoviruses. Generated viruses were collected 2-3 days later from both the supernatant and adherent cells by three freeze-thaw cycles. HEK293 cells were seeded in 96 well plate and next day, the virus suspension was diluted at 1:100. Later, 10μl virus supernatant was added to cells in 90μl culture media. To calculate virions, further 10 fold serial dilutions were done. One week later, cytopathic effect (CPE) was tested based on cell morphology.

Alternatively, viral titres were determined using immunofluorescence using antibodies against hexon protein. Infected cells were visualized using an EVOS microscope after 3X PBSP wash. To test the effect of E2F1 on hC9ORF78, 0.25×10^5^ WI-38 cells were plated in 6cm dishes in 10%FBS DMEM medium. Next day, cells were serum starved and incubated for 3 days. Viruses at indicated MOI were added. Cells were washed with PBS after overnight incubation with viruses, followed by addition of starvation medium. Cells were harvested for western blotting analysis 48 h after adding viruses.

## ABBREVIATIONS

ACA: Anti-centromeric antibody
SAC: Spindle assembly checkpoint
APC/C: Anaphase promoting complex/cyclosome
BSA: Bovine serum albumin
CENP-A: centromeric protein A
CPC: Chromosome Passenger Complex
INCENP: Inner centromere protein
MCC: mitotic checkpoint complex
PBS: Phosphate buffered saline
hC9ORF78: human Chromosome 9 Open Reading Frame 78
TERF2IP: Telomeric Repeat binding Factor 2 Interacting protein
POT1: Protection Of Telomeres 1
PRPF8: Pre-mRNA processing Factor 8

## Acknowledgements

The authors thank Tomer Avidor-Reiss for help with microscopy, Song-Tao Liu for help in kinetochore staining methodology, and Nishanth Kuganesan for help in adenovirus propagation.

## Competing interests

The authors have no competing interests to declare.

## Funding

This project was supported by grant R15GM120712 to W.R.T.

## Data availability

This study generated no large-scale datasets.

## Notes

### Competing Interest Statement

The authors have declared no competing interest.

